## Supplementary material for "Light-gated Integrator for Highlighting Kinase Activity in Living Cells": KINACT Supplementary Figures

**Supplementary Figure 1**

**Supplementary Figure 2**

**Supplementary Figure 3**

**Supplementary Figure 4**

**Supplementary Figure 5**

**Supplementary Figure 6**

**Supplementary Figure 7**

**Supplementary Figure 8**

**Supplementary Figure 9**

**Supplementary Figure 10**

**Supplementary Figure 11**

**Supplementary Figure 12**

**Supplementary Figure 13**

**Supplementary Figure 14**

**Supplementary Figure 15**

**Supplementary Figure 16**

**Sequences of constructs**

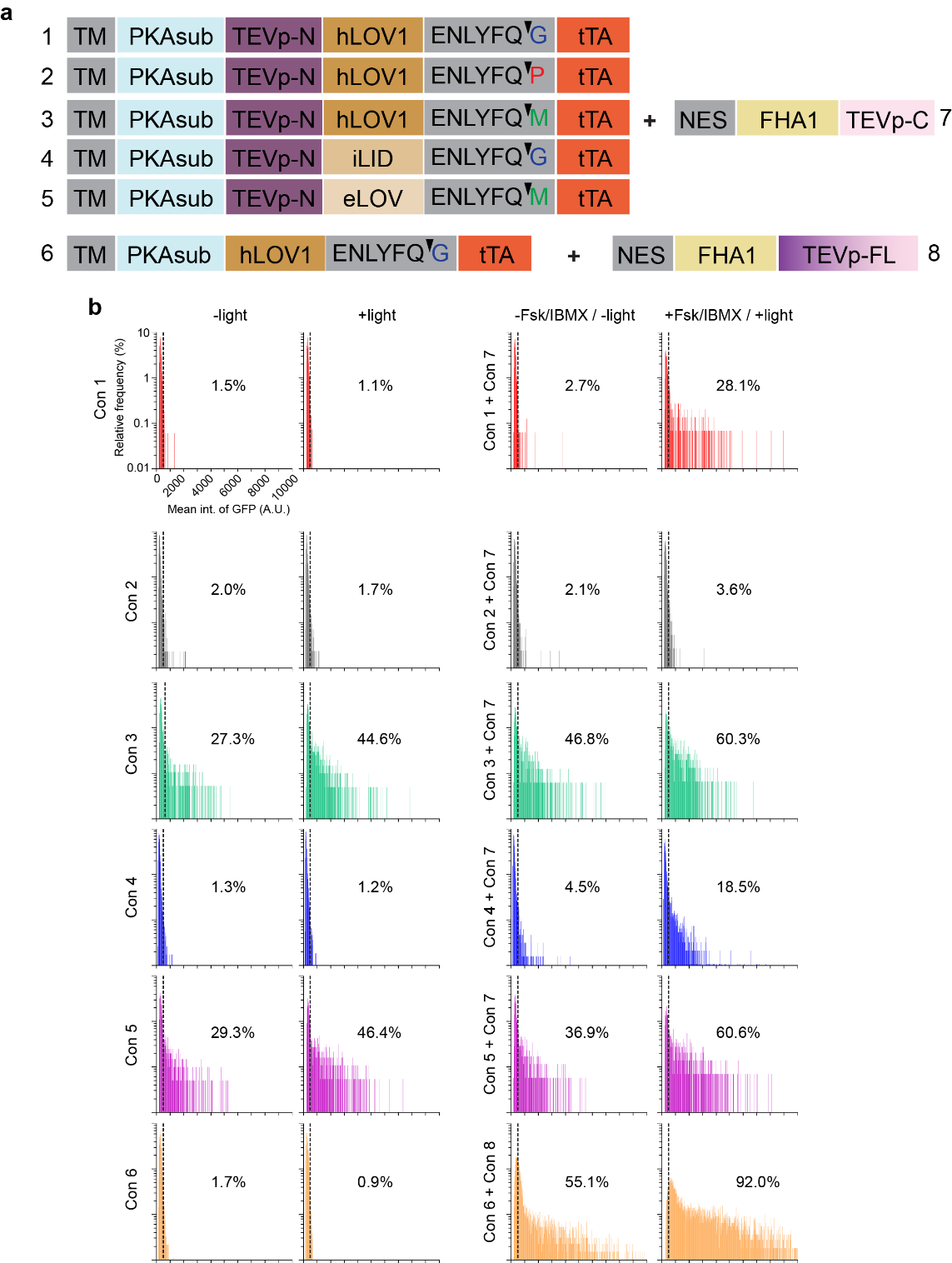

**Supplementary Fig. 1 | KINACT optimization. a**, Domain structures of various combinations of A-KINACT components 1 and 2. **b**, Histograms showing induced H2B-EGFP expression in HEK293T cells expressing different versions of component 1 alone or co-expressed with different versions of component 2 under dark or light conditions. The fraction of H2B-EGFP^+^ cells above the threshold cut-off (dashed line) is indicated. Data from 3 independent experiments.

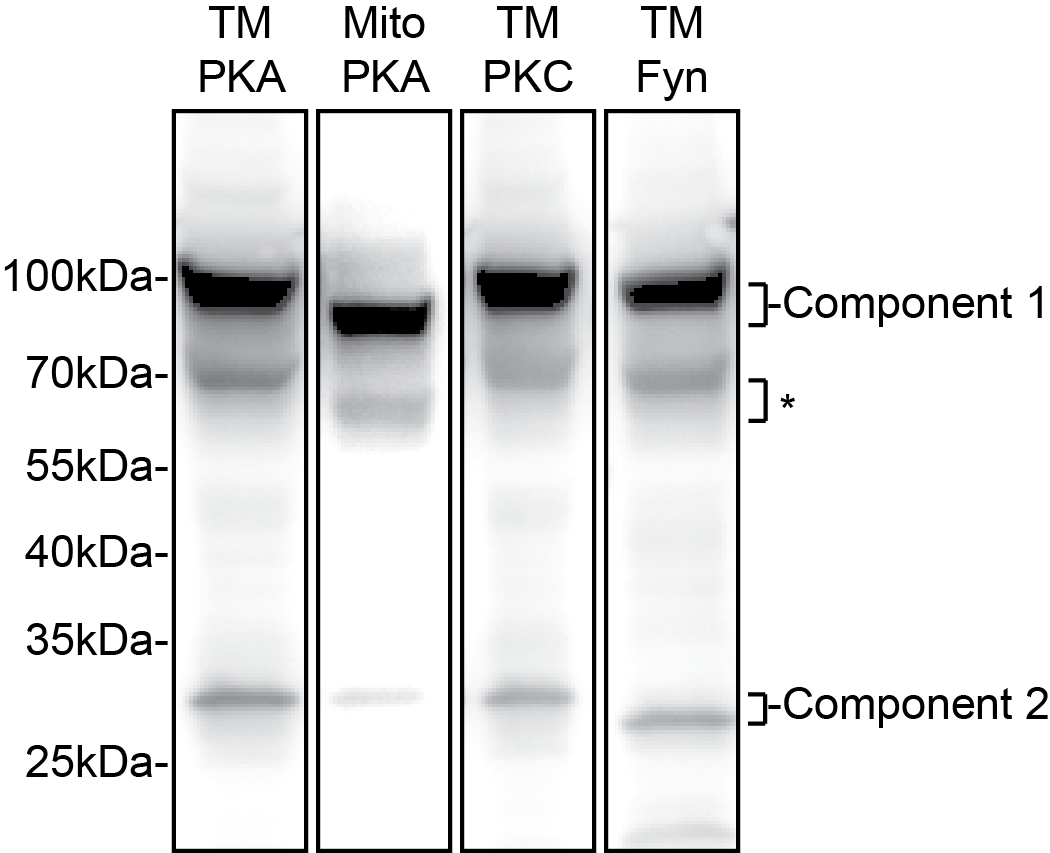

**Supplementary Fig. 2 | Representative western blot characterization of main KINACT integrators expression.** Bands at ~100 kDa correspond to component 1 of each KINACT, and bands between 25 kDa and 35 kDa correspond to component 2 of each KINACT. Both components 1 and 2 were fused to an N-terminal Myc epitope and blotted using an anti-Myc antibody. *Weak bands at ~70kDa correspond to truncated component 1 lacking the tTA domain, which has no transcriptional activity.

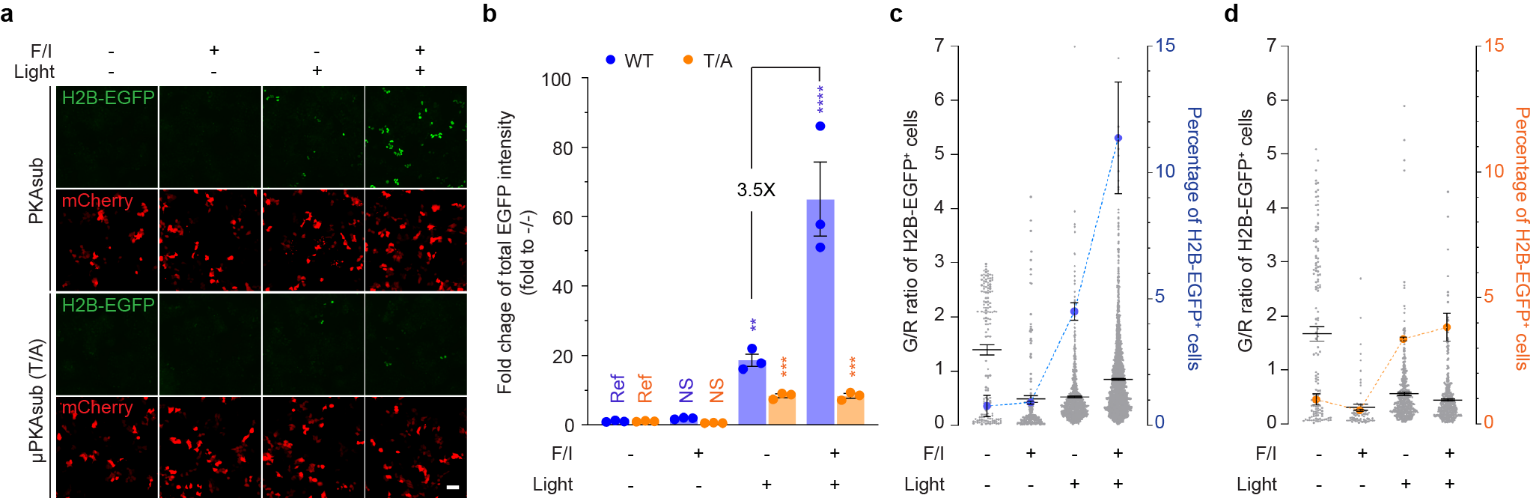

**Supplementary Fig. 3 | Transient expression of A-KINACT for cumulative PKA activity in TetO-H2B-EGFP HEK293T cells. a**, Snapshot imaging of cumulative PKA activity in live cells. Four treatment conditions were applied: -F/I/-light, +F/I/-light, -F/I/+light and +F/I/+light. Scale bar, 10 μm. **b**, Statistical quantification of total EGFP intensity under all conditions. *P* = 0.0011 (WT, -F/I/+light), *P* < 0.0001 (WT, +F/I/+light), *P* = 0.0002 (T/A, +F/I/-light) and *P* = 0.0002 (T/A, -F/I/+light). NS, not significant. Statistical analysis was performed using ordinary one-way ANOVA followed by Dunnett’s multiple-comparison test. **c**, The fraction and G/R ratio of H2B-EGFP^+^ cells expressing A-KINACT (WT). n = 143, n = 183, n = 836 and n = 2081 cells. **d**, The fraction and G/R ratio of H2B-EGFP^+^ cells expressing A-KINACT (T/A). n = 133, n = 67, n = 442 and n = 452 cells. For **b**-**d**, data from 3 independent experiments. Data are mean ± s.e.m.

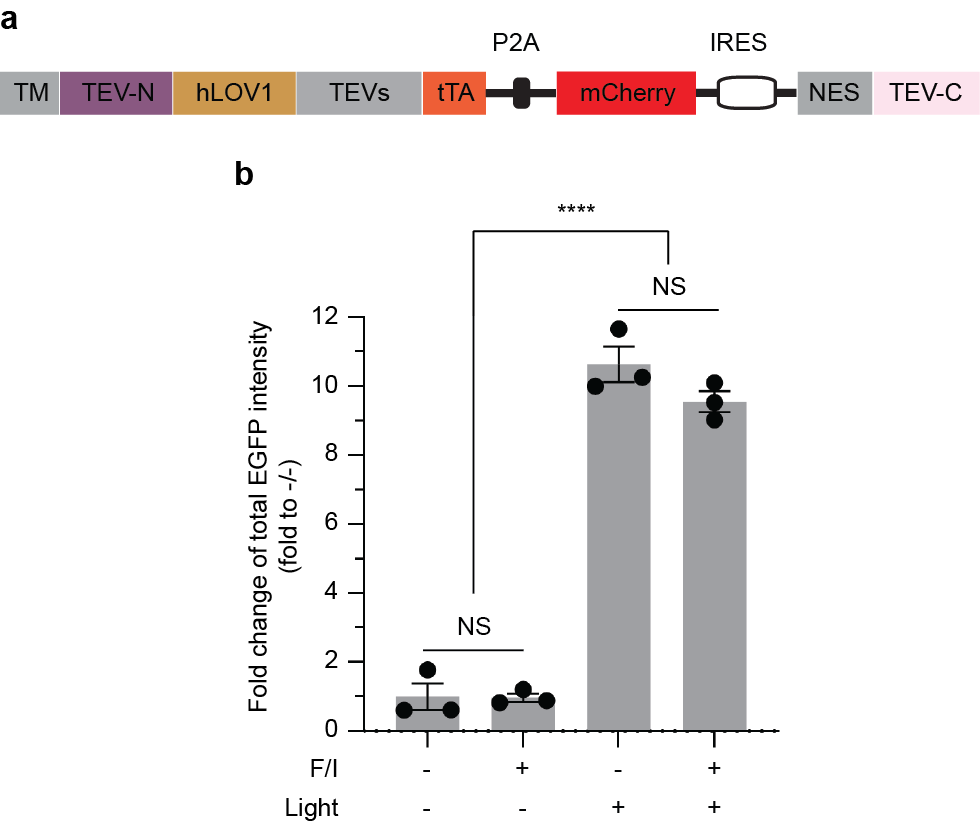

**Supplementary Fig. 4 | Overexpression of TEV-only integrator in HEK293T cells indicating spontaneous binding of components 1 and 2. a**, Domain structure of TEV-only integrator. The PKA substrate peptide and FHA1 domain were removed from A-KINACT. **k**, Statistical quantification of total EGFP intensity under -F/I/-light, +F/I/-light, -F/I/+light and +F/I/+light conditions. *****P* < 0.0001. n = 3 independent experiments. Statistical analysis was performed using ordinary one-way ANOVA followed by Tukey’s multiple-comparison test. NS, not significant. Data are mean ± s.e.m.

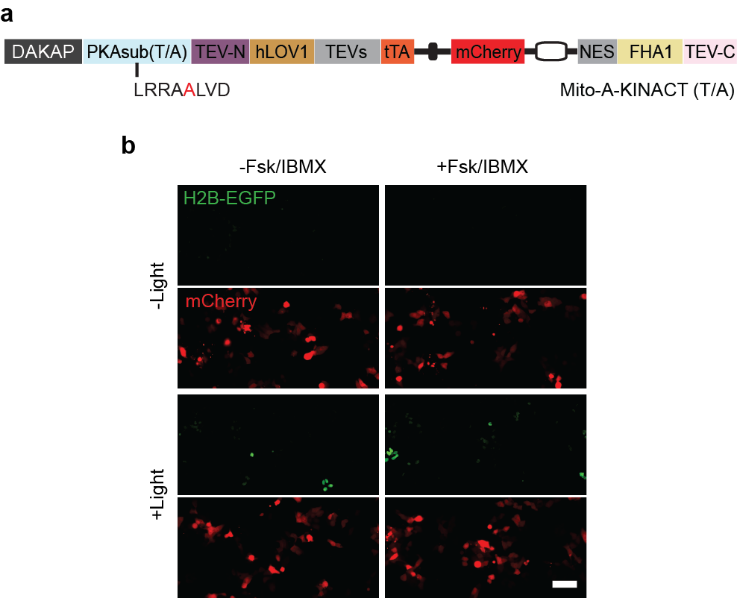

**Supplementary Fig. 5** **| Images of compartmentalized PKA activity using nonphosphorylatable Mito-targeted A-KINACT (T/A). a**, Domain structure of outer mitochondrial membrane-targeted A-KINACT (T/A). **b**, Snapshot imaging of spontaneously induced H2B-EGFP expression. Four treatment conditions were used: -F/I/-light, +F/I/-light, -F/I/+light and +F/I/+light. Scale bar, 10 μm.

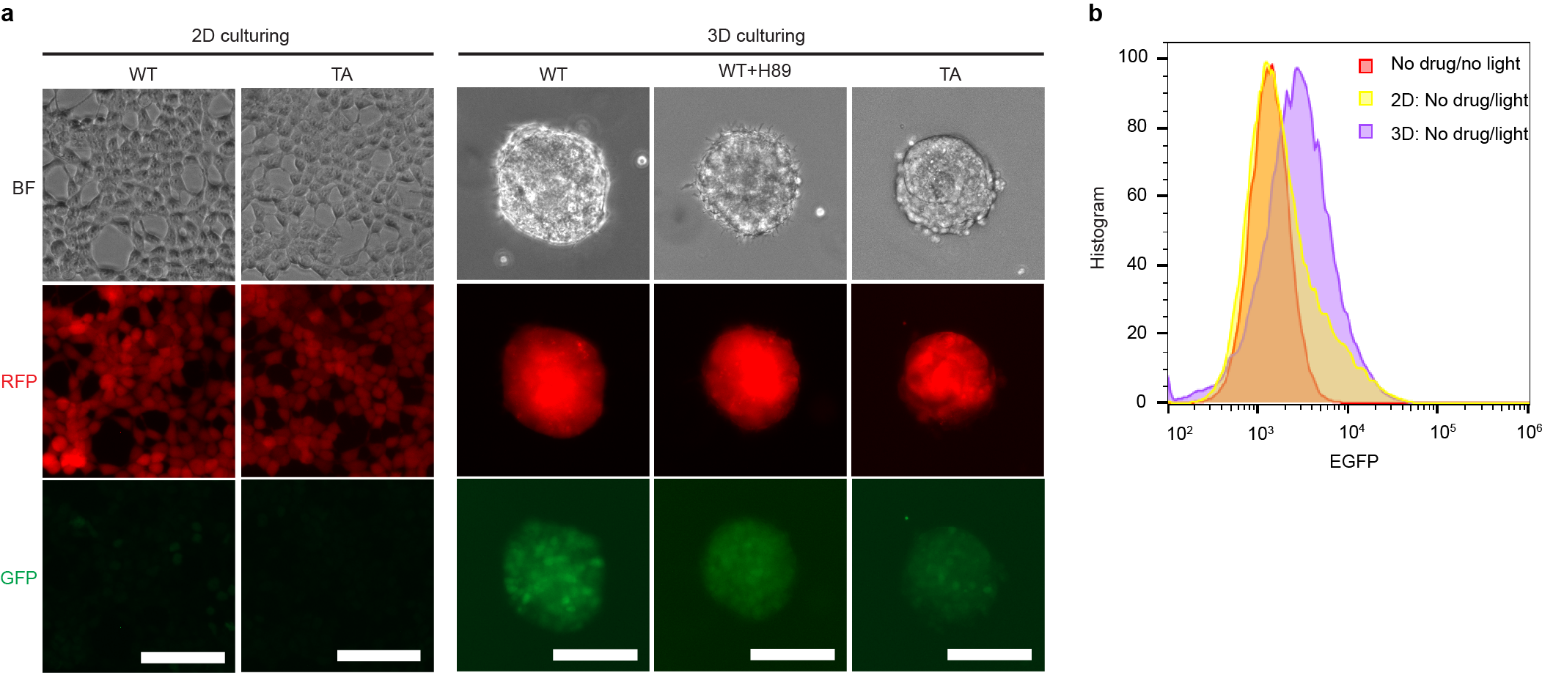

**Supplementary Fig. 6** **| Comparison of basal PKA activities between 2D and 3D culture conditions. a**, Epifluorescence imaging of blue light-induced H2B-EGFP expression in A-KINACT and A-KINACT (T/A) dual stable cells performed on an EVOS tissue culture microscope (Invitrogen). Scale bar, 50 μm. **b**, Histogram of EGFP intensity in A-KINACT dual stable cells by flow-cytometry.

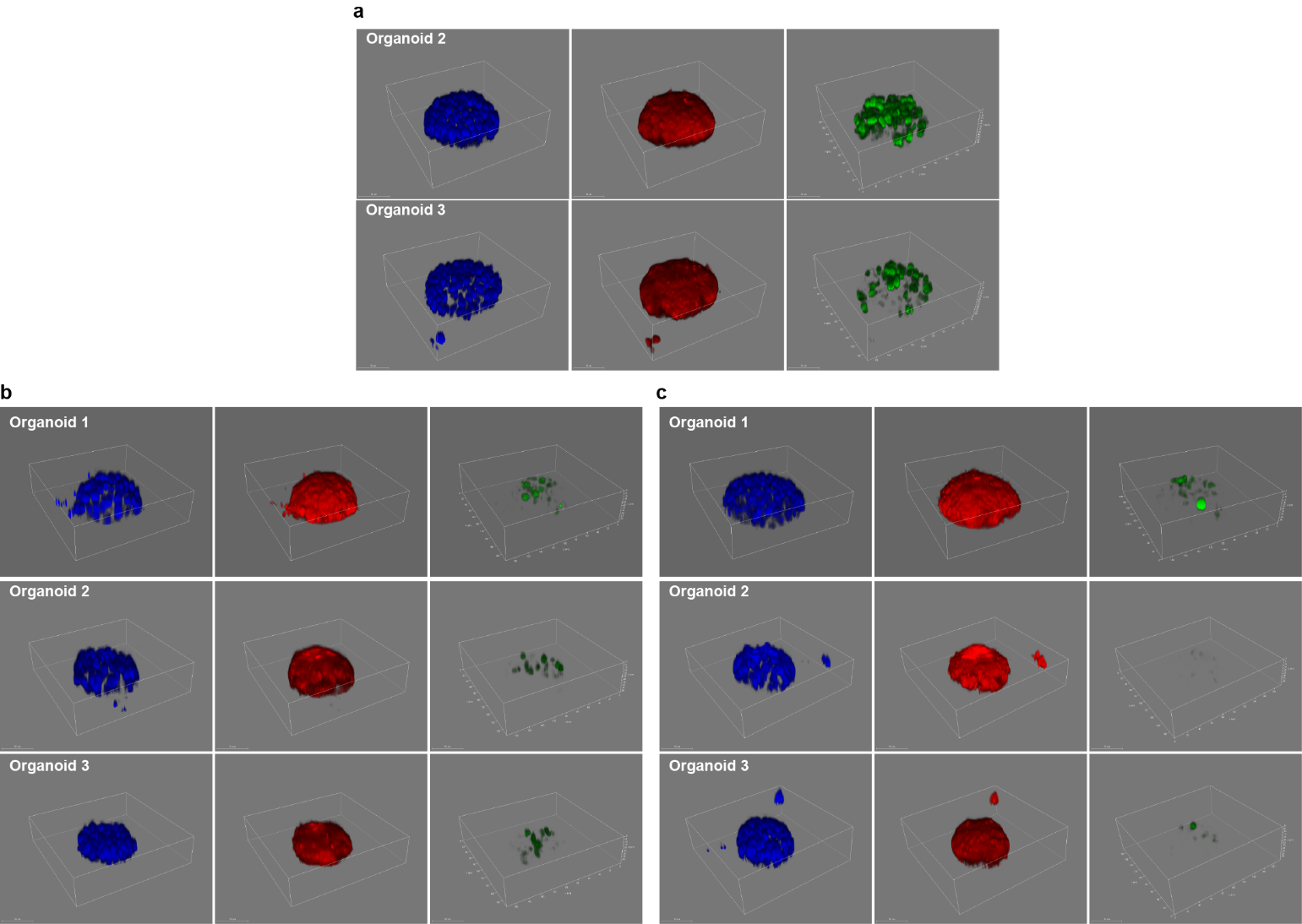

**Supplementary Fig. 7** **| Representative 3D imaging of heterogeneous basal PKA activity within HEK293T organoids. a**, Two replicates of 3D PKA activity imaging in A-KINACT organoids. **b**, Three replicates of 3D PKA activity imaging in A-KINACT (WT) organoids pretreated with H89. **c**, Three replicates of 3D PKA activity imaging in A-KINACT (T/A) organoids.

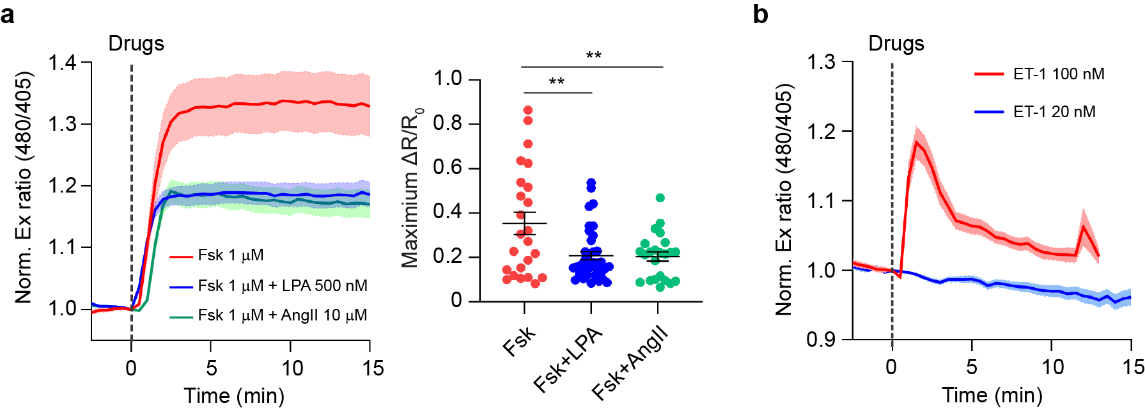

**Supplementary Fig. 8 | Validation of PKA activity responding to Gα_i_- and Gα_q_-linked drugs using ExRai-AKAR2. a**, Average time courses (left) and maximum responses (ΔR/R_0_) of the PKA activty stimulated with 1 μM Fsk only (red, n = 24 cells from 2 independent experiments), premixed 1 μM Fsk and 500 nM LPA (blue, n = 38 cells from 3 independent experiments) and premixed 1 μM Fsk and 10 μM Ang II (green, n = 23 cells from 1 independent experiment). **b**, Average time courses of the PKA response stimulated with 20 nM ET-1 (blue curve, n = 8 cells) and 100 nM ET-1 (red curve, n = 8 cells). Data from 1 independent experiment. The solid lines represent the mean; shaded areas, s.e.m. Dashed lines indicate drug addition.

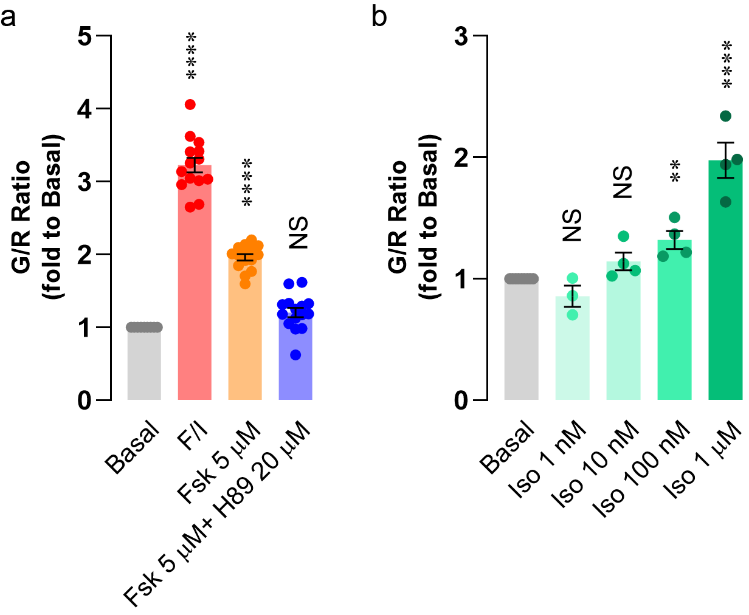

**Supplementary Fig. 9 | Testing the sensitivity of the A-KINACT readout on a multi-well plate reader. a**, A-KINACT induction of H2B-EGFP expression in response to no stimulation (Basal, n = 6 wells), and stimulation with F/I (50 μM/100 μM, positive control, n = 14 wells), low-dose Fsk (5 μM, n = 15 wells) and Fsk (5 μM) with H89 (20 μM) pretreatment (n = 15 wells). *****P* < 0.0001. **b**, Iso dose response of A-KINACT. Basal (n = 6 wells), Iso (1 nM, n = 3 wells), Iso (10 nM, n = 4 wells), Iso (100 nM, *P* = 0.008, n = 4 wells) and Iso (1 μM, *P* < 0.0001, n = 4 wells). Statistical analysis was performed using ordinary one-way ANOVA followed by Dunnett’s multiple-comparison test. NS, not significant. Data are mean ± s.e.m.

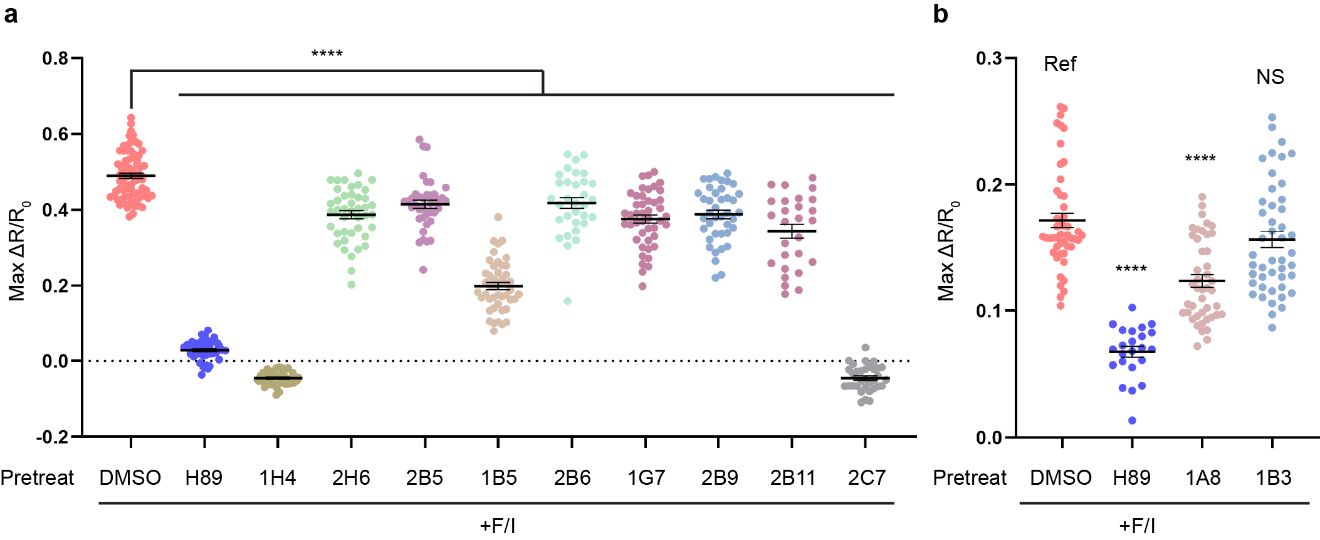

**Supplementary Fig. 10** **| Validation of inhibitory effects of 11 kinase inhibitors on PKA activity. a**, Maximum responses (ΔR/R_0_) of GR-AKARev in HEK293T cells pretreated with each inhibitor (10 μM) and then treatment with F/I (50 μM/100 μM). DMSO, n = 73 cells from 5 independent experiments. H89, n = 43 cells; 1H4, n = 39 cells; 2H6, n = 42 cells; 2B5, n = 39 cells; 1B5, n = 46 cells; 2B6, n = 32 cells; 1G7, n = 46 cells; 2B9, n = 40 cells; 2B11, n = 27 cells; 2C7, n = 33 cells; data from 3 independent experiments. **b**, Maximum responses (ΔR/R_0_) of Booster-PKA in HEK293T cells pretreated with each inhibitors (10 μM) and then treated with F/I (50 μM/100 μM). DMSO, n = 48 cells from 4 independent experiments; H89, n = 23 cells from 2 independent experiments; 1A8, n = 44 cells from 3 independent experiments; 1B3, n = 46 cells from 3 independent experiments. Statistical analysis was performed using ordinary one-way ANOVA followed by Dunnett’s multiple-comparison test. ****P<0.0001. NS, not significant. Data are mean ± s.e.m.

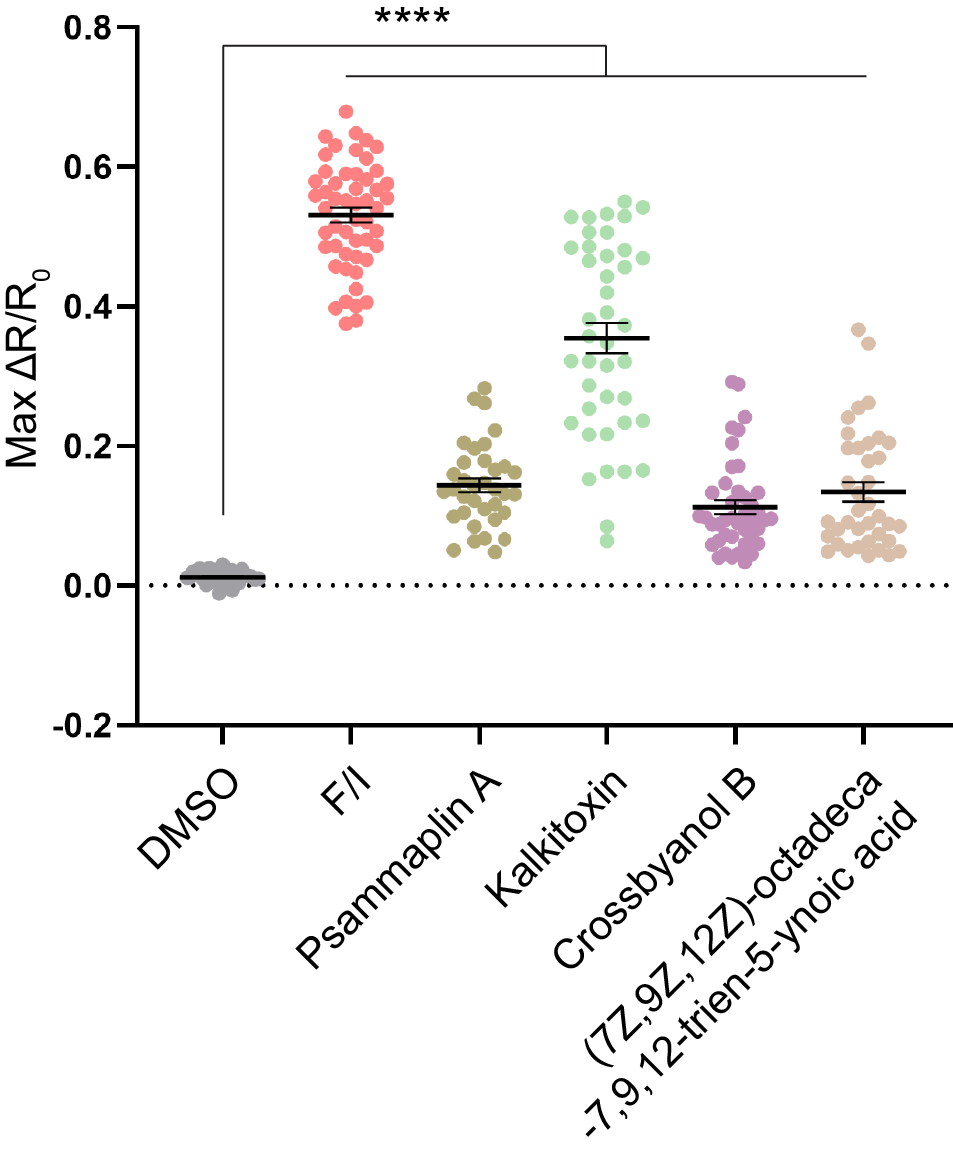

**Supplementary Fig. 11 | Validation of 4 PKA activators from marine natural product library.** Maximum responses (ΔR/R_0_) of AKAR3ev in HEK293T cells treated with each compound (2 μg ml^−1^). F/I (50 μM/100 μM) treatment shown as a positive control. DMSO, n = 35 cells; F/I, n = 52 cells; Psa-A, n = 35 cells; Kal, n = 41 cells; Cro-B, n = 42 cells; 7Z, n = 39 cells. All data from 3 independent experiments. Statistical analysis was performed using ordinary one-way ANOVA followed by Dunnett’s multiple-comparison test. *****P* < 0.0001. Data are mean ± s.e.m.

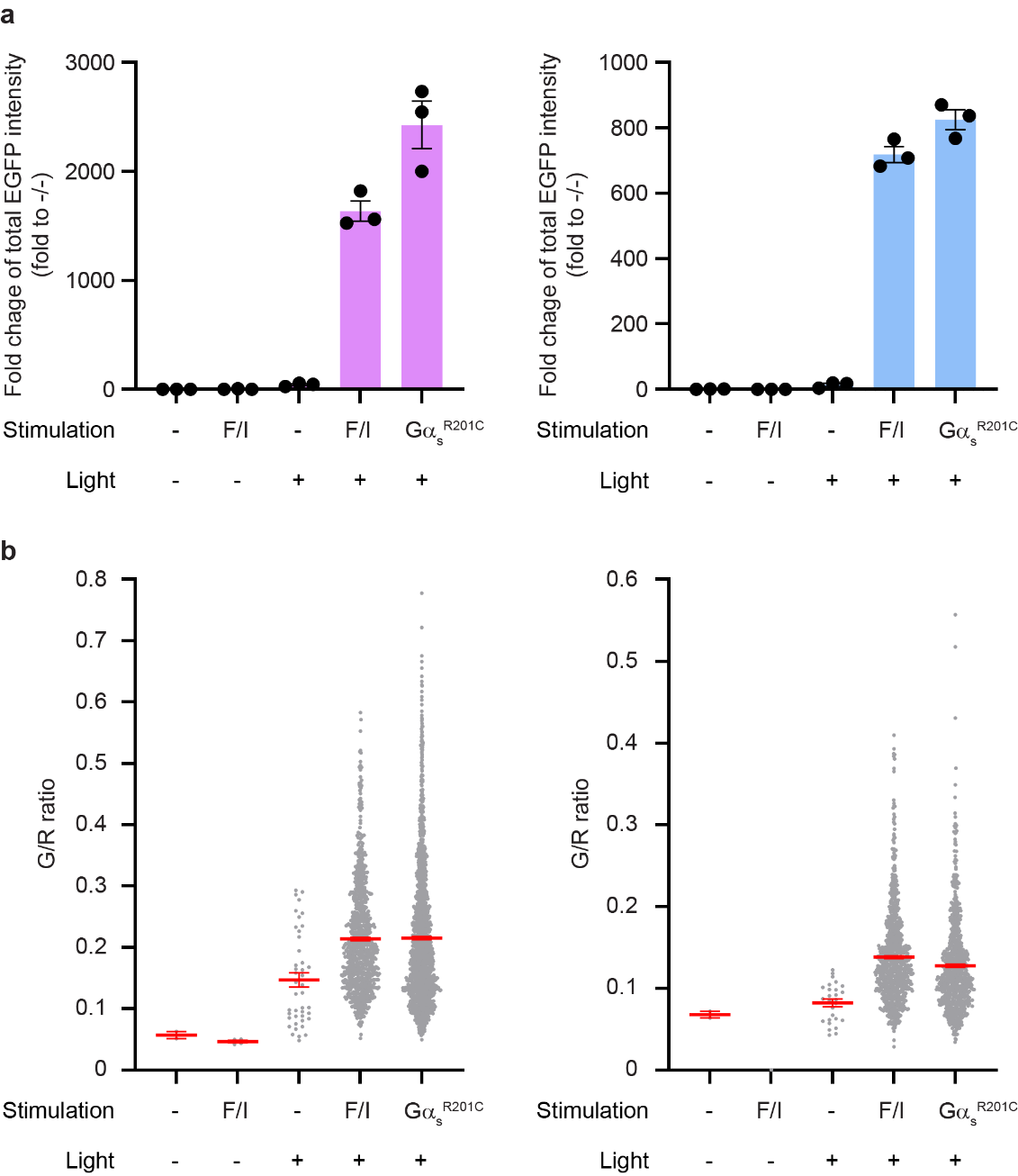

**Supplementary Fig. 12** **|** **Characterization of A-KINACT effector (PKI-EGFP) and A-KINACT control (EGFP) dual-stable cell lines.** **a**, Statistical quantification of total EGFP intensity in A-KINACT effector cells (left) and A-KINACT control cells (right) under -F/I/-light, +F/I/-light, -F/I/+light, +F/I/+light and +Gα_s_^R201C^/+light conditions. n = 3 independent experiments. NS, not significant. Statistical analysis was performed using ordinary one-way ANOVA followed by Dunnett’s multiple-comparison test. **b**, G/R ratio of individual PKI-EGFP^+^ cells (left, n = 2, n = 4, n = 43, n = 964 and n = 1913 cells from 3 independent experiments.) and EGFP^+^ cells (right, n = 2, n = 0, n = 27, n = 973 and n = 854 cells from 3 independent experiments.). Data are mean ± s.e.m.

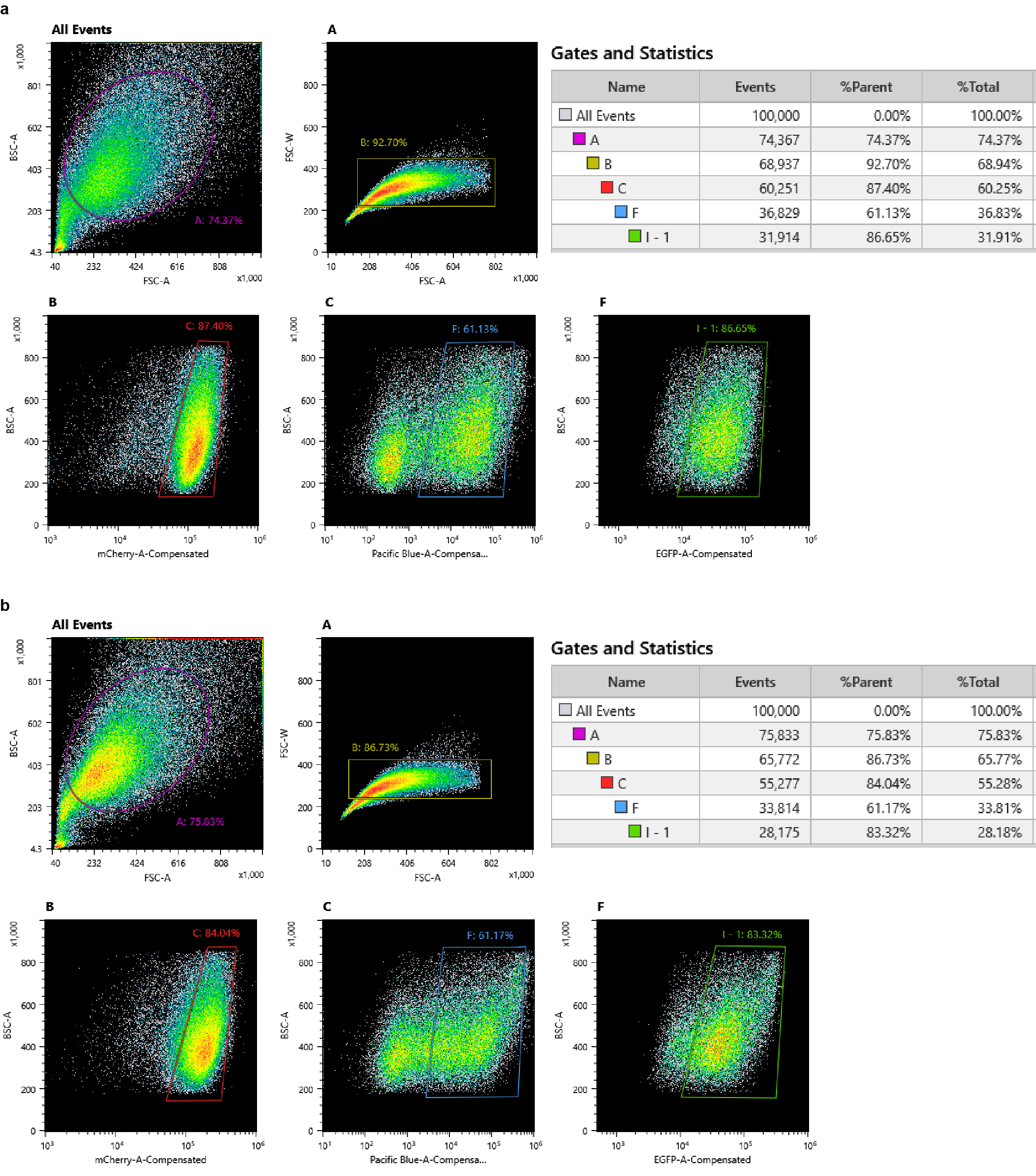
**Supplementary Fig. 13** **| Procedure of triple-color FACS for sorting PKI-EGFP^+^ cells (a) and EGFP^+^ cells (b).** Gate A removing cell debris. Gate B indicating single cells. Gate C indicating mCherry positive cells. Gate F indicating mTagBFP2-Gα_s_^R201C^ positive cells. Gate I-1 indicating H2B-EGFP positive cells which were finally collected.

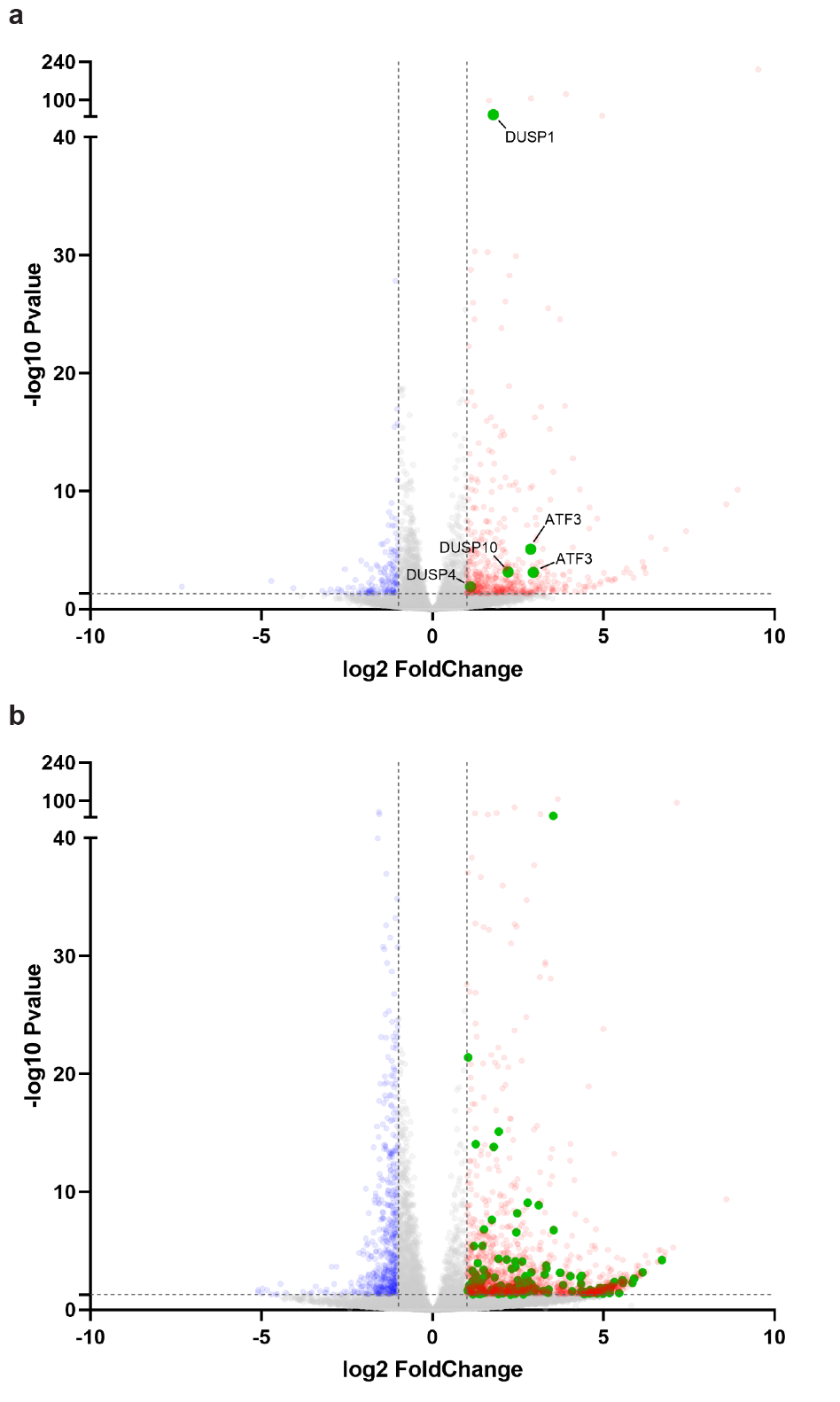

**Supplementary Fig. 14** **| Distribution of key PKA signatures and ERK signatures. a**, Volcano plot showing relative enrichment of transcripts in A-KINACT control (EGFP) cells under Gα_s_^R201C^ /light condition versus no Gα_s_^R201C^ /no light condition. Transcripts corresponding to 4 genes associated with Negative regulation of ERK cascade pathway are highlighted (green points). **b**, Volcano plot showing relative enrichment of transcripts in A-KINACT effector (PKI-EGFP) cells under Gα_s_^R201C^/light condition versus no Gα_s_^R201C^ /no light condition. Transcripts corresponding to 111 genes associated with Cell cycle pathway (sharing 60 genes associated with Mitotic cell cycle proc. pathway; 68 genes associated with Mitotic cell cycle pathway; 87 genes associated with Cell cycle proc. pathway) are highlighted (green points). Cut-off: P value<0.05; Fold change>2. See Supplementary Table 5 and 6 for gene lists.

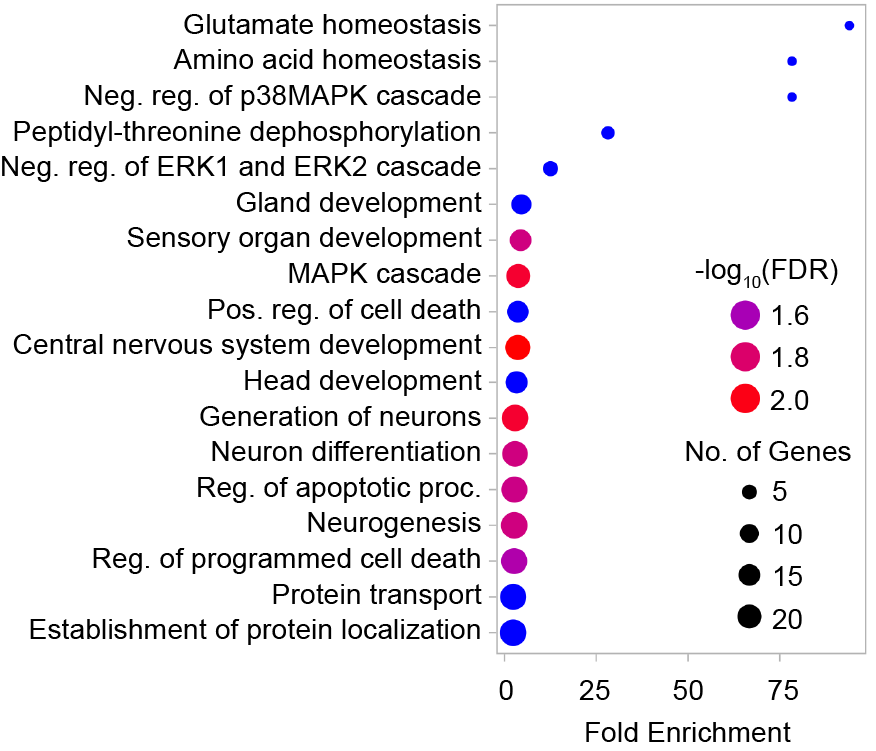

**Supplementary Fig. 15** **| GO analysis of 106 PKA-upregulated genes in A-KINACT control cells.**

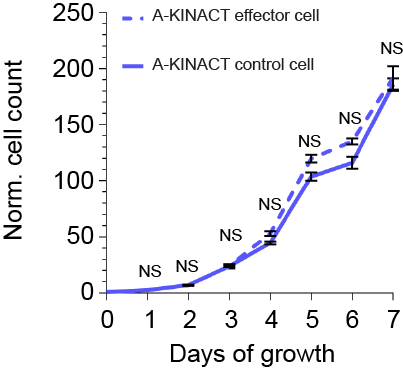

**Supplementary Fig. 16 |** **Growth curves of A-KINACT effector (PKI-EGFP) cells and A-KINACT control (EGFP) cells without Gα_s_^R201C^ overexpression and blue light illumination.** There was no significant difference at each time point. n = 3 independent experiments. Statistical analysis was performed using unpaired *t* test. Data are mean ± s.e.m.

**Sequences of constructs**

**A-KINACT Construct 1**

**METDTLLLWVLLLWVPGSTGDEQKLISEEDLNAVGQDTQEVIVVPHSLPFKVVVISAILALVVLTIISLIILIMLWQKKPR**GGSGGLEGM**LRRATLVD**ACGGGGSGGGGSGGGGRSGGSM**LQLPPLERLTLEMGESLFKGPRDYNPISSTICHLTNESDGHTTSLYGIGFGPFIITNKHLFRRNNGTLLVQSLHGVFKVKNTTTLQQHLIDGRDMIIIRMPKDFPPFPQKLKFREPQREERICLVTTNFQ**ELAEKLAGLDINGGASG**SRATTLERIEKSFVITDPRLPDNPIIFVSDSFLQLTEYSREEILGRNCRFLQGPETDRATVRKIRDAIDNQTEVTVQLINYTKSGKKFWNVFHLQPMRDYKGDVQYFIGVQLDGTERLHGAAEREAVCLVKKTAFQIAENLYFQGSRLDKSKVINSALELLNEVGIEGLTTRKLAQKLGVEQPTLYWHVKNKRALLDALAIEMLDRHHTHFCPLEGESWQDFLRNNAKSFRCALLSHRDGAKVHLGTRPTEKQYETLENQLAFLCQQGFSLENALYALSAVGHFTLGCVLEDQEHQVAKEERETPTTDSMPPLLRQAIELFDHQGAEPAFLFGLELIICGLEKQLKCESGSAYSRARTKNNYGSTIEGLLDLPDDDAPEEAGLAAPRLSFLPAGHTRRLSTAPPTDVSLGDELHLDGEDVAMAHADALDDFDLDMLGDGDSPGPGFTPHDSAPYGALDMADFEFEQMFTDALGIDEYGG***

**Ig Kappa secretory element: 1-21 a.a.**

**Myc-epitope: 22-31 a.a.**

**PDGFRβ transmembrane helix: 32-81 a.a.**

**PKAsub: 91-98 a.a.**

**nuclear export signal (NES): 121-132 a.a.**

**TEVp-N: 133-250 a.a.**

**hLOV1: 268-405 a.a.**

**TEVseq-G: 406-412 a.a.**

**tTA-VP16: 413-746 a.a.**

**A-KINACT Construct 2**

**METDTLLLWVLLLWVPGSTGDEQKLISEEDLNAVGQDTQEVIVVPHSLPFKVVVISAILALVVLTIISLIILIMLWQKKPR**GGSGGLEGM**LRRATLVD**ACGGGGSGGGGSGGGGRSGGSM**LQLPPLERLTLEMGESLFKGPRDYNPISSTICHLTNESDGHTTSLYGIGFGPFIITNKHLFRRNNGTLLVQSLHGVFKVKNTTTLQQHLIDGRDMIIIRMPKDFPPFPQKLKFREPQREERICLVTTNFQ**ELAEKLAGLDINGGASG**SRATTLERIEKSFVITDPRLPDNPIIFVSDSFLQLTEYSREEILGRNCRFLQGPETDRATVRKIRDAIDNQTEVTVQLINYTKSGKKFWNVFHLQPMRDYKGDVQYFIGVQLDGTERLHGAAEREAVCLVKKTAFQIAENLYFQPSRLDKSKVINSALELLNEVGIEGLTTRKLAQKLGVEQPTLYWHVKNKRALLDALAIEMLDRHHTHFCPLEGESWQDFLRNNAKSFRCALLSHRDGAKVHLGTRPTEKQYETLENQLAFLCQQGFSLENALYALSAVGHFTLGCVLEDQEHQVAKEERETPTTDSMPPLLRQAIELFDHQGAEPAFLFGLELIICGLEKQLKCESGSAYSRARTKNNYGSTIEGLLDLPDDDAPEEAGLAAPRLSFLPAGHTRRLSTAPPTDVSLGDELHLDGEDVAMAHADALDDFDLDMLGDGDSPGPGFTPHDSAPYGALDMADFEFEQMFTDALGIDEYGG***

**Ig Kappa secretory element: 1-21 a.a.**

**Myc-epitope: 22-31 a.a.**

**PDGFRβ transmembrane helix: 32-81 a.a.**

**PKAsub: 91-98 a.a.**

**nuclear export signal (NES): 121-132 a.a.**

**TEVp-N: 133-250 a.a.**

**hLOV1: 268-405 a.a.**

**TEVseq-P: 406-412 a.a.**

**tTA-VP16: 413-746 a.a.**

**A-KINACT Construct 3**

**METDTLLLWVLLLWVPGSTGDEQKLISEEDLNAVGQDTQEVIVVPHSLPFKVVVISAILALVVLTIISLIILIMLWQKKPR**GGSGGLEGM**LRRATLVD**ACGGGGSGGGGSGGGGRSGGSM**LQLPPLERLTLEMGESLFKGPRDYNPISSTICHLTNESDGHTTSLYGIGFGPFIITNKHLFRRNNGTLLVQSLHGVFKVKNTTTLQQHLIDGRDMIIIRMPKDFPPFPQKLKFREPQREERICLVTTNFQ**ELAEKLAGLDINGGASG**SRATTLERIEKSFVITDPRLPDNPIIFVSDSFLQLTEYSREEILGRNCRFLQGPETDRATVRKIRDAIDNQTEVTVQLINYTKSGKKFWNVFHLQPMRDYKGDVQYFIGVQLDGTERLHGAAEREAVCLVKKTAFQIAENLYFQMSRLDKSKVINSALELLNEVGIEGLTTRKLAQKLGVEQPTLYWHVKNKRALLDALAIEMLDRHHTHFCPLEGESWQDFLRNNAKSFRCALLSHRDGAKVHLGTRPTEKQYETLENQLAFLCQQGFSLENALYALSAVGHFTLGCVLEDQEHQVAKEERETPTTDSMPPLLRQAIELFDHQGAEPAFLFGLELIICGLEKQLKCESGSAYSRARTKNNYGSTIEGLLDLPDDDAPEEAGLAAPRLSFLPAGHTRRLSTAPPTDVSLGDELHLDGEDVAMAHADALDDFDLDMLGDGDSPGPGFTPHDSAPYGALDMADFEFEQMFTDALGIDEYGG***

**Ig Kappa secretory element: 1-21 a.a.**

**Myc-epitope: 22-31 a.a.**

**PDGFRβ transmembrane helix: 32-81 a.a.**

**PKAsub: 91-98 a.a.**

**nuclear export signal (NES): 121-132 a.a.**

**TEVp-N: 133-250 a.a.**

**hLOV1: 268-405 a.a.**

**TEVseq-M: 406-412 a.a.**

**tTA-VP16: 413-746 a.a.**

**A-KINACT Construct 4**

**METDTLLLWVLLLWVPGSTGDEQKLISEEDLNAVGQDTQEVIVVPHSLPFKVVVISAILALVVLTIISLIILIMLWQKKPR**GGSGGLEGM**LRRATLVD**ACGGGGSGGGGSGGGGRSGGSM**LQLPPLERLTLEMGESLFKGPRDYNPISSTICHLTNESDGHTTSLYGIGFGPFIITNKHLFRRNNGTLLVQSLHGVFKVKNTTTLQQHLIDGRDMIIIRMPKDFPPFPQKLKFREPQREERICLVTTNFQ**ELGSGSG**EFLATTLERIEKNFVITDPRLPDNPIIFASDSFLQLTEYSREEILGRNCRFLQGPETDRATVRKIRDAIDNQTEVTVQLINYTKSGKKFWNVFHLQPMRDYKGDVQYFIGVQLDGTERLHGAAEREAVCLIKKTAFQIAENLYFQGSRLDKSKVINSALELLNEVGIEGLTTRKLAQKLGVEQPTLYWHVKNKRALLDALAIEMLDRHHTHFCPLEGESWQDFLRNNAKSFRCALLSHRDGAKVHLGTRPTEKQYETLENQLAFLCQQGFSLENALYALSAVGHFTLGCVLEDQEHQVAKEERETPTTDSMPPLLRQAIELFDHQGAEPAFLFGLELIICGLEKQLKCESGSAYSRARTKNNYGSTIEGLLDLPDDDAPEEAGLAAPRLSFLPAGHTRRLSTAPPTDVSLGDELHLDGEDVAMAHADALDDFDLDMLGDGDSPGPGFTPHDSAPYGALDMADFEFEQMFTDALGIDEYGG***

**Ig Kappa secretory element: 1-21 a.a.**

**Myc-epitope: 22-31 a.a.**

**PDGFRβ transmembrane helix: 32-81 a.a.**

**PKAsub: 91-98 a.a.**

**nuclear export signal (NES): 121-132 a.a.**

**TEVp-N: 133-250 a.a.**

**iLID: 258-396 a.a.**

**TEVseq-G: 397-403 a.a.**

**tTA-VP16: 404-737 a.a.**

**A-KINACT Construct 5**

**METDTLLLWVLLLWVPGSTGDEQKLISEEDLNAVGQDTQEVIVVPHSLPFKVVVISAILALVVLTIISLIILIMLWQKKPR**GGSGGLEGM**LRRATLVD**ACGGGGSGGGGSGGGGRSGGSM**LQLPPLERLTLEMGESLFKGPRDYNPISSTICHLTNESDGHTTSLYGIGFGPFIITNKHLFRRNNGTLLVQSLHGVFKVKNTTTLQQHLIDGRDMIIIRMPKDFPPFPQKLKFREPQREERICLVTTNFQ**ELAEKLAGLDINGGASG**SRATTLERIEKSFVITDPRLPDNPIIFVSDSFLQLTEYSREEILGRNCRFLQGPETDRATVRKIRDAIDNQTEVTVQLINYTKSGKKFWNLFHLQPMRDQKGDVQYFIGVQLDGTERVRDAAEREAVMLVKKTAEEIDEAAKENLYFQMGGGSDYKDDDDKSRLDKSKVINSALELLNEVGIEGLTTRKLAQKLGVEQPTLYWHVKNKRALLDALAIEMLDRHHTHFCPLEGESWQDFLRNNAKSFRCALLSHRDGAKVHLGTRPTEKQYETLENQLAFLCQQGFSLENALYALSAVGHFTLGCVLEDQEHQVAKEERETPTTDSMPPLLRQAIELFDHQGAEPAFLFGLELIICGLEKQLKCESGSAYSRARTKNNYGSTIEGLLDLPDDDAPEEAGLAAPRLSFLPAGHTRRLSTAPPTDVSLGDELHLDGEDVAMAHADALDDFDLDMLGDGDSPGPGFTPHDSAPYGALDMADFEFEQMFTDALGIDEYGG***

**Ig Kappa secretory element: 1-21 a.a.**

**Myc-epitope: 22-31 a.a.**

**PDGFRβ transmembrane helix: 32-81 a.a.**

**PKAsub: 91-98 a.a.**

**nuclear export signal (NES): 121-132 a.a.**

**TEVp-N: 133-250 a.a.**

**hLOV1: 268-409 a.a.**

**TEVseq-M: 410-416 a.a.**

**Flag-tTA-VP16: 421-762 a.a.**

**A-KINACT Construct 6**

**METDTLLLWVLLLWVPGSTGDEQKLISEEDLNAVGQDTQEVIVVPHSLPFKVVVISAILALVVLTIISLIILIMLWQKKPR**GGSGGLEGM**LRRATLVD**ACGGGGSGGGGSGGGGRSGGSM**LQLPPLERLTLE**ELAEKLAGLDINGGASG**SRATTLERIEKSFVITDPRLPDNPIIFVSDSFLQLTEYSREEILGRNCRFLQGPETDRATVRKIRDAIDNQTEVTVQLINYTKSGKKFWNVFHLQPMRDYKGDVQYFIGVQLDGTERLHGAAEREAVCLVKKTAFQIAENLYFQGSRLDKSKVINSALELLNEVGIEGLTTRKLAQKLGVEQPTLYWHVKNKRALLDALAIEMLDRHHTHFCPLEGESWQDFLRNNAKSFRCALLSHRDGAKVHLGTRPTEKQYETLENQLAFLCQQGFSLENALYALSAVGHFTLGCVLEDQEHQVAKEERETPTTDSMPPLLRQAIELFDHQGAEPAFLFGLELIICGLEKQLKCESGSAYSRARTKNNYGSTIEGLLDLPDDDAPEEAGLAAPRLSFLPAGHTRRLSTAPPTDVSLGDELHLDGEDVAMAHADALDDFDLDMLGDGDSPGPGFTPHDSAPYGALDMADFEFEQMFTDALGIDEYGG***

**Ig Kappa secretory element: 1-21 a.a.**

**Myc-epitope: 22-31 a.a.**

**PDGFRβ transmembrane helix: 32-81 a.a.**

**PKAsub: 91-98 a.a.**

**nuclear export signal (NES): 121-132 a.a.**

**hLOV1: 150-287 a.a.**

**TEVseq-G: 288-294 a.a.**

**tTA-VP16: 295-628 a.a.**

**A-KINACT Construct 7**

M**EQKLISEEDLLQLPPLERLTLE**AT**MKFSQEQIGENIVCRVICTTGQIPIRDLSADISQVLKEKRSIKKVWTFGRNPACDYHLGNISRLSNKHFQILLGEDGNLLLNDISTNGTWLNGQKVEKNSNQLLSQGDEITVGVGVESDILSLVIFINDKFKQCLEQNKVDR**LQELGGGGRSGGGGS**TKSMSSMVSDTSCTFPSSDGIFWKHWIQTKDGQCGSPLVSTRDGFIVGIHSASNFTNTNNYFTSVPKNFMELLTNQEAQQWVSGWRLNADSVLWGGHKVFMV***

**Myc-epitope: 2-11 a.a.**

**nuclear export signal (NES): 12-23 a.a.**

**FHA1: 26-167 a.a.**

**TEVp-C: 183-284 a.a.**

**A-KINACT Construct 8**

M**EQKLISEEDLLQLPPLERLTLE**AT**MKFSQEQIGENIVCRVICTTGQIPIRDLSADISQVLKEKRSIKKVWTFGRNPACDYHLGNISRLSNKHFQILLGEDGNLLLNDISTNGTWLNGQKVEKNSNQLLSQGDEITVGVGVESDILSLVIFINDKFKQCLEQNKVDR**LQELGGGGRSGGGGS**MGESLFKGPRDYNPISSTICHLTNESDGHTTSLYGIGFGPFIITNKHLFRRNNGTLLVQSLHGVFKVKNTTTLQQHLIDGRDMIIIRMPKDFPPFPQKLKFREPQREERICLVTTNFQTKSMSSMVSDTSCTFPSSDGIFWKHWIQTKDGQCGSPLVSTRDGFIVGIHSASNFTNTNNYFTSVPKNFMELLTNQEAQQWVSGWRLNADSVLWGGHKVFMV***

**Myc-epitope: 2-11 a.a.**

**nuclear export signal (NES): 12-23 a.a.**

**FHA1: 26-167 a.a.**

**TEVp: 183-402 a.a.**

**Mito-A-KINACT**

**MAIQLRSLFPLALPGMLALLGWWWFFSRKKEQKLISEEDL**GGSGGLEGM**LRRATLVD**ACGGGGSGGGGSGGGGRSGGSM**LQLPPLERLTLEMGESLFKGPRDYNPISSTICHLTNESDGHTTSLYGIGFGPFIITNKHLFRRNNGTLLVQSLHGVFKVKNTTTLQQHLIDGRDMIIIRMPKDFPPFPQKLKFREPQREERICLVTTNFQ**ELAEKLAGLDINGGASG**SRATTLERIEKSFVITDPRLPDNPIIFVSDSFLQLTEYSREEILGRNCRFLQGPETDRATVRKIRDAIDNQTEVTVQLINYTKSGKKFWNVFHLQPMRDYKGDVQYFIGVQLDGTERLHGAAEREAVCLVKKTAFQIAENLYFQGSRLDKSKVINSALELLNEVGIEGLTTRKLAQKLGVEQPTLYWHVKNKRALLDALAIEMLDRHHTHFCPLEGESWQDFLRNNAKSFRCALLSHRDGAKVHLGTRPTEKQYETLENQLAFLCQQGFSLENALYALSAVGHFTLGCVLEDQEHQVAKEERETPTTDSMPPLLRQAIELFDHQGAEPAFLFGLELIICGLEKQLKCESGSAYSRARTKNNYGSTIEGLLDLPDDDAPEEAGLAAPRLSFLPAGHTRRLSTAPPTDVSLGDELHLDGEDVAMAHADALDDFDLDMLGDGDSPGPGFTPHDSAPYGALDMADFEFEQMFTDALGIDEYGG***

**DAKAP1-tag: 1-30 a.a.**

**Myc-epitope: 31-40 a.a.**

**PKAsub: 50-57 a.a.**

**nuclear export signal (NES): 80-91 a.a.**

**TEVp-N: 92-209 a.a.**

**hLOV1: 227-364 a.a.**

**TEVseq-G: 365-371 a.a.**

**tTA-VP16: 372-705 a.a.**

**C-KINACT Component 1**

**METDTLLLWVLLLWVPGSTGDEQKLISEEDLNAVGQDTQEVIVVPHSLPFKVVVISAILALVVLTIISLIILIMLWQKKPR**GGSGGLEGM**RFRRFQTLKDKAKA**ACGGGGSGGGGSGGGGRSGGSM**LQLPPLERLTLEMGESLFKGPRDYNPISSTICHLTNESDGHTTSLYGIGFGPFIITNKHLFRRNNGTLLVQSLHGVFKVKNTTTLQQHLIDGRDMIIIRMPKDFPPFPQKLKFREPQREERICLVTTNFQ**ELAEKLAGLDINGGASG**SRATTLERIEKSFVITDPRLPDNPIIFVSDSFLQLTEYSREEILGRNCRFLQGPETDRATVRKIRDAIDNQTEVTVQLINYTKSGKKFWNVFHLQPMRDYKGDVQYFIGVQLDGTERLHGAAEREAVCLVKKTAFQIAENLYFQGSRLDKSKVINSALELLNEVGIEGLTTRKLAQKLGVEQPTLYWHVKNKRALLDALAIEMLDRHHTHFCPLEGESWQDFLRNNAKSFRCALLSHRDGAKVHLGTRPTEKQYETLENQLAFLCQQGFSLENALYALSAVGHFTLGCVLEDQEHQVAKEERETPTTDSMPPLLRQAIELFDHQGAEPAFLFGLELIICGLEKQLKCESGSAYSRARTKNNYGSTIEGLLDLPDDDAPEEAGLAAPRLSFLPAGHTRRLSTAPPTDVSLGDELHLDGEDVAMAHADALDDFDLDMLGDGDSPGPGFTPHDSAPYGALDMADFEFEQMFTDALGIDEYGG***

**Ig Kappa secretory element: 1-21 a.a.**

**Myc-epitope: 22-31 a.a.**

**PDGFRβ transmembrane helix: 32-81 a.a.**

**PKCsub: 91-104 a.a.**

**nuclear export signal (NES): 127-138 a.a.**

**TEVp-N: 139-256 a.a.**

**hLOV1: 274-411 a.a.**

**TEVseq-G: 412-418 a.a.**

**tTA-VP16: 419-752 a.a.**

**C-KINACT Component 2**

Same with A-KINACT construct 6

**F-KINACT Component 1**

**METDTLLLWVLLLWVPGSTGDEQKLISEEDLNAVGQDTQEVIVVPHSLPFKVVVISAILALVVLTIISLIILIMLWQKKPR**GGSGGLEGM**EKIEGTYGVV**ACGGGGSGGGGSGGGGRSGGSM**LQLPPLERLTLEMGESLFKGPRDYNPISSTICHLTNESDGHTTSLYGIGFGPFIITNKHLFRRNNGTLLVQSLHGVFKVKNTTTLQQHLIDGRDMIIIRMPKDFPPFPQKLKFREPQREERICLVTTNFQ**ELAEKLAGLDINGGASG**SRATTLERIEKSFVITDPRLPDNPIIFVSDSFLQLTEYSREEILGRNCRFLQGPETDRATVRKIRDAIDNQTEVTVQLINYTKSGKKFWNVFHLQPMRDYKGDVQYFIGVQLDGTERLHGAAEREAVCLVKKTAFQIAENLYFQGSRLDKSKVINSALELLNEVGIEGLTTRKLAQKLGVEQPTLYWHVKNKRALLDALAIEMLDRHHTHFCPLEGESWQDFLRNNAKSFRCALLSHRDGAKVHLGTRPTEKQYETLENQLAFLCQQGFSLENALYALSAVGHFTLGCVLEDQEHQVAKEERETPTTDSMPPLLRQAIELFDHQGAEPAFLFGLELIICGLEKQLKCESGSAYSRARTKNNYGSTIEGLLDLPDDDAPEEAGLAAPRLSFLPAGHTRRLSTAPPTDVSLGDELHLDGEDVAMAHADALDDFDLDMLGDGDSPGPGFTPHDSAPYGALDMADFEFEQMFTDALGIDEYGG***

**Ig Kappa secretory element: 1-21 a.a.**

**Myc-epitope: 22-31 a.a.**

**PDGFRβ transmembrane helix: 32-81 a.a.**

**Fynsub: 91-100 a.a.**

**nuclear export signal (NES): 123-134 a.a.**

**TEVp-N: 135-252 a.a.**

**hLOV1: 270-407 a.a.**

**TEVseq-G: 408-414 a.a.**

**tTA-VP16: 415-748 a.a.**

**F-KINACT Component 2**

M**EQKLISEEDLLQLPPLERLTLE**AT**MWYFGKITRRESERLLLNPENPRGTFLVRESETTKGAYALSVSDFDNAKGLNVKHYKIRKLDSGGFYITSRTQFSSLQQLVAYYSKHADGLCHRLTNV**LQELGGGGRSGGGGS**TKSMSSMVSDTSCTFPSSDGIFWKHWIQTKDGQCGSPLVSTRDGFIVGIHSASNFTNTNNYFTSVPKNFMELLTNQEAQQWVSGWRLNADSVLWGGHKVFMV***

**Myc-epitope: 2-11 a.a.**

**nuclear export signal (NES): 12-23 a.a.**

**SH2(C185A): 26-123 a.a.**

**TEVp-C: 139-240 a.a.**

TetO-reporters

TetO-H2B-EGFP

cgagtttaccactccctatcagtgatagagaaaagtgaaagtcgagtttaccactccctatcagtgatagagaaaagtgaaagtcgagtttaccactccctatcagtgatagagaaaagtgaaagtcgagtttaccactccctatcagtgatagagaaaagtgaaagtcgagtttaccactccctatcagtgatagagaaaagtgaaagtcgagtttaccactccctatcagtgatagagaaaagtgaaagtcgagtttaccactccctatcagtgatagagaaaagtgaaagtcgagtttaccactccctatcagtgatagagaaaagtgaaagtcgagtttaccactccctatcagtgatagagaaaagtgaaagtcgagtttaccactccctatcagtgatagagaaaagtgaaagtcgagtttaccactccctatcagtgatagagaaaagtgaaagtcgagtttaccactccctatcagtgatagagaaaagtgaaagtcgagtttaccactccctatcagtgatagagaaaagtgaaagtcgagctcggtacgctatggcatgcatgtgtcgcgacctgcaggccctgaagttcatctgcaccaccggcaagctgcccgtgccctggcccaccctcgtgaccaccctgacctggggcgtgcagtgcttcgcccgctaccccgaccacatgaagcagcacgacttcttcaagtccgccatgcccgaaggctacgtccaggagcgcaccatcttcttcaaggacgacggcaactacaagacccgcgccgaggtgaagttcgagggcgacaccctggtgaaccgcatcgagctgaagggcatcgacttcaaggaggacggcaacatcctggggcacaagctggagtacaacgccatcagcgacaacgtctatatcaccgccgacaagcagaagaacggcatcaaggccaacttcaagagctagccctatataagcagagctcgtttagtgaaccgtcagatcgcctggagacgccatccacgctgttttgacctccatagaagacaccgggaccgatccagcctccgcggccccggtaccgaattcaaggcctctcgagcctctagaaggtggcggaATGCCAGAGCCAGCGAAGTCTGCTCCCGCCCCGAAAAAGGGCTCCAAGAAGGCGGTGACTAAGGCGCAGAAGAAAGGCGGCAAGAAGCGCAAGCGCAGCCGCAAGGAGAGCTATTCCATCTATGTGTACAAGGTTCTGAAGCAGGTCCACCCTGACACCGGCATTTCGTCCAAGGCCATGGGCATCATGAATTCGTTTGTGAACGACATTTTCGAGCGCATCGCAGGTGAGGCTTCCCGCCTGGCGCATTACAACAAGCGCTCGACCATCACCTCCAGGGAGATCCAGACGGCCGTGCGCCTGCTGCTGCCTGGGGAGTTGGCCAAGCACGCCGTGTCCGAGGGTACTAAGGCCATCACCAAGTACACCAGCGCTAAGgatccaccggtcgccaccATGGTGAGCAAGGGCGAGGAGCTGTTCACCGGGGTGGTGCCCATCCTGGTCGAGCTGGACGGCGACGTAAACGGCCACAAGTTCAGCGTGTCCGGCGAGGGCGAGGGCGATGCCACCTACGGCAAGCTGACCCTGAAGTTCATCTGCACCACCGGCAAGCTGCCCGTGCCCTGGCCCACCCTCGTGACCACCCTGACCTACGGCGTGCAGTGCTTCAGCCGCTACCCCGACCACATGAAGCAGCACGACTTCTTCAAGTCCGCCATGCCCGAAGGCTACGTCCAGGAGCGCACCATCTTCTTCAAGGACGACGGCAACTACAAGACCCGCGCCGAGGTGAAGTTCGAGGGCGACACCCTGGTGAACCGCATCGAGCTGAAGGGCATCGACTTCAAGGAGGACGGCAACATCCTGGGGCACAAGCTGGAGTACAACTACAACAGCCACAACGTCTATATCATGGCCGACAAGCAGAAGAACGGCATCAAGGTGAACTTCAAGATCCGCCACAACATCGAGGACGGCAGCGTGCAGCTCGCCGACCACTACCAGCAGAACACCCCCATCGGCGACGGCCCCGTGCTGCTGCCCGACAACCACTACCTGAGCACCCAGTCCGCCCTGAGCAAAGACCCCAACGAGAAGCGCGATCACATGGTCCTGCTGGAGTTCGTGACCGCCGCCGGGATCACTCTCGGCATGGACGAGCTGTACAAGTAG

TetO-PKI-EGFP

cgagtttaccactccctatcagtgatagagaaaagtgaaagtcgagtttaccactccctatcagtgatagagaaaagtgaaagtcgagtttaccactccctatcagtgatagagaaaagtgaaagtcgagtttaccactccctatcagtgatagagaaaagtgaaagtcgagtttaccactccctatcagtgatagagaaaagtgaaagtcgagtttaccactccctatcagtgatagagaaaagtgaaagtcgagtttaccactccctatcagtgatagagaaaagtgaaagtcgagtttaccactccctatcagtgatagagaaaagtgaaagtcgagtttaccactccctatcagtgatagagaaaagtgaaagtcgagtttaccactccctatcagtgatagagaaaagtgaaagtcgagtttaccactccctatcagtgatagagaaaagtgaaagtcgagtttaccactccctatcagtgatagagaaaagtgaaagtcgagtttaccactccctatcagtgatagagaaaagtgaaagtcgagctcggtacgctatggcatgcatgtgtcgcgacctgcaggccctgaagttcatctgcaccaccggcaagctgcccgtgccctggcccaccctcgtgaccaccctgacctggggcgtgcagtgcttcgcccgctaccccgaccacatgaagcagcacgacttcttcaagtccgccatgcccgaaggctacgtccaggagcgcaccatcttcttcaaggacgacggcaactacaagacccgcgccgaggtgaagttcgagggcgacaccctggtgaaccgcatcgagctgaagggcatcgacttcaaggaggacggcaacatcctggggcacaagctggagtacaacgccatcagcgacaacgtctatatcaccgccgacaagcagaagaacggcatcaaggccaacttcaagagctagccctatataagcagagctcgtttagtgaaccgtcagatcgcctggagacgccatccacgctgttttgacctccatagaagacaccgggaccgatccagcctccgcggccccggtaccgaattcaaggcctctcgagcctctagaaggtggcggaATGACATATGCAGATTTTATTGCTTCAGGAAGAACAGGTAGAAGAAATGCAATATGGgatccaccggtcgccaccATGGTGAGCAAGGGCGAGGAGCTGTTCACCGGGGTGGTGCCCATCCTGGTCGAGCTGGACGGCGACGTAAACGGCCACAAGTTCAGCGTGTCCGGCGAGGGCGAGGGCGATGCCACCTACGGCAAGCTGACCCTGAAGTTCATCTGCACCACCGGCAAGCTGCCCGTGCCCTGGCCCACCCTCGTGACCACCCTGACCTACGGCGTGCAGTGCTTCAGCCGCTACCCCGACCACATGAAGCAGCACGACTTCTTCAAGTCCGCCATGCCCGAAGGCTACGTCCAGGAGCGCACCATCTTCTTCAAGGACGACGGCAACTACAAGACCCGCGCCGAGGTGAAGTTCGAGGGCGACACCCTGGTGAACCGCATCGAGCTGAAGGGCATCGACTTCAAGGAGGACGGCAACATCCTGGGGCACAAGCTGGAGTACAACTACAACAGCCACAACGTCTATATCATGGCCGACAAGCAGAAGAACGGCATCAAGGTGAACTTCAAGATCCGCCACAACATCGAGGACGGCAGCGTGCAGCTCGCCGACCACTACCAGCAGAACACCCCCATCGGCGACGGCCCCGTGCTGCTGCCCGACAACCACTACCTGAGCACCCAGTCCGCCCTGAGCAAAGACCCCAACGAGAAGCGCGATCACATGGTCCTGCTGGAGTTCGTGACCGCCGCCGGGATCACTCTCGGCATGGACGAGCTGTACAAGTAG

TetO-EGFP

cgagtttaccactccctatcagtgatagagaaaagtgaaagtcgagtttaccactccctatcagtgatagagaaaagtgaaagtcgagtttaccactccctatcagtgatagagaaaagtgaaagtcgagtttaccactccctatcagtgatagagaaaagtgaaagtcgagtttaccactccctatcagtgatagagaaaagtgaaagtcgagtttaccactccctatcagtgatagagaaaagtgaaagtcgagtttaccactccctatcagtgatagagaaaagtgaaagtcgagtttaccactccctatcagtgatagagaaaagtgaaagtcgagtttaccactccctatcagtgatagagaaaagtgaaagtcgagtttaccactccctatcagtgatagagaaaagtgaaagtcgagtttaccactccctatcagtgatagagaaaagtgaaagtcgagtttaccactccctatcagtgatagagaaaagtgaaagtcgagtttaccactccctatcagtgatagagaaaagtgaaagtcgagctcggtacgctatggcatgcatgtgtcgcgacctgcaggccctgaagttcatctgcaccaccggcaagctgcccgtgccctggcccaccctcgtgaccaccctgacctggggcgtgcagtgcttcgcccgctaccccgaccacatgaagcagcacgacttcttcaagtccgccatgcccgaaggctacgtccaggagcgcaccatcttcttcaaggacgacggcaactacaagacccgcgccgaggtgaagttcgagggcgacaccctggtgaaccgcatcgagctgaagggcatcgacttcaaggaggacggcaacatcctggggcacaagctggagtacaacgccatcagcgacaacgtctatatcaccgccgacaagcagaagaacggcatcaaggccaacttcaagagctagccctatataagcagagctcgtttagtgaaccgtcagatcgcctggagacgccatccacgctgttttgacctccatagaagacaccgggaccgatccagcctccgcggccccggtaccgaattcaaggcctctcgagcctctagaaggtggcggaATGGTGAGCAAGGGCGAGGAGCTGTTCACCGGGGTGGTGCCCATCCTGGTCGAGCTGGACGGCGACGTAAACGGCCACAAGTTCAGCGTGTCCGGCGAGGGCGAGGGCGATGCCACCTACGGCAAGCTGACCCTGAAGTTCATCTGCACCACCGGCAAGCTGCCCGTGCCCTGGCCCACCCTCGTGACCACCCTGACCTACGGCGTGCAGTGCTTCAGCCGCTACCCCGACCACATGAAGCAGCACGACTTCTTCAAGTCCGCCATGCCCGAAGGCTACGTCCAGGAGCGCACCATCTTCTTCAAGGACGACGGCAACTACAAGACCCGCGCCGAGGTGAAGTTCGAGGGCGACACCCTGGTGAACCGCATCGAGCTGAAGGGCATCGACTTCAAGGAGGACGGCAACATCCTGGGGCACAAGCTGGAGTACAACTACAACAGCCACAACGTCTATATCATGGCCGACAAGCAGAAGAACGGCATCAAGGTGAACTTCAAGATCCGCCACAACATCGAGGACGGCAGCGTGCAGCTCGCCGACCACTACCAGCAGAACACCCCCATCGGCGACGGCCCCGTGCTGCTGCCCGACAACCACTACCTGAGCACCCAGTCCGCCCTGAGCAAAGACCCCAACGAGAAGCGCGATCACATGGTCCTGCTGGAGTTCGTGACCGCCGCCGGGATCACTCTCGGCATGGACGAGCTGTACAAGTAG
